# Cholinergic stimulation modulates the functional composition of CA3 cell types in the hippocampus

**DOI:** 10.1101/2022.04.04.486716

**Authors:** Christopher Jon Puhl, Winnie Wefelmeyer, Juan Burrone

## Abstract

The functional heterogeneity of hippocampal CA3 pyramidal neurons has emerged as a key aspect of circuit function. Here, we explored the effects of long-term cholinergic activity on the functional heterogeneity of CA3 pyramidal neurons in organotypic slices. Application of agonists to either acetylcholine receptors (AChRs) generally, or muscarinic AChRs (mAChRs) specifically, induced robust increases in network activity in the low-gamma range. Prolonged AChR stimulation for 48 hrs uncovered a population of hyperadapting CA3 pyramidal neurons that typically fired a single, early action potential in response to current injection. Although these neurons were present in control networks, their proportions were dramatically increased following long-term cholinergic activity. Characterised by the presence of a strong M-current, the hyperadaptation phenotype was abolished by acute application of either M-channel antagonists or the re-application of AChR agonists. We conclude that long-term mAChR activation modulates the intrinsic excitability of a subset of CA3 pyramidal cells, uncovering a highly plastic cohort of neurons that are sensitive to chronic ACh modulation. Our findings provide evidence for the activity-dependent plasticity of functional heterogeneity in the hippocampus.

## Introduction

Neurons in the brain are highly heterogeneous, making their classification complicated (Zeng and Sanes, 2017). Aside from neuronal morphology or genetic profiles, one of the strongest features that define neuronal cell types is their function (Sharpee, 2014). At its most basic, the intrinsic excitability of neurons – the ability to produce an output to a given input – provides a first order description of cell function and is key to understanding circuit function. To date, it remains unclear whether the proportions of functionally-defined neurons is plastic, allowing the repertoire of neuron types in the brain to be modulated.

In contrast to GABAergic interneurons (Lim et al., 2018), pyramidal cells in the hippocampus have typically been thought of as a homogeneous population of cells, at least within the specific area they populate – from CA1 to CA3 subfields. However, closer inspections of their functional properties have gradually uncovered important differences even within this supposedly uniform neuronal population (Strange et al., 2014). In CA1, for example, pyramidal cells mapped across different spatial axes were shown to have distinct functional properties (Cavalieri et al., 2021; Milior et al., 2016; Slomianka et al., 2011). Recently, two clearly distinct populations of CA3 pyramidal neurons have been described, with markedly different functional and morphological characteristics (Hunt et al., 2018). Whereas the majority of CA3 pyramidal cells showed non-adapting spike outputs, a subset of neurons were characterised by their high levels of adaptation and transient high frequency firing. The latter population of neurons lacked inputs from the dentate gyrus, had distinct dendritic morphologies and were differentially modulated by ACh (Hunt et al., 2018). However, what determines the functional differences between these two populations of CA3 neurons and their modulation by activity remains unexplored.

The neurotransmitter Acetylcholine (ACh) is thought to play a fundamental role in the process of learning and memory in the hippocampus (Hasselmo, 2006; Solari and Hangya, 2018). Cholinergic medial septum neurons project their axons to the hippocampus where they control network activity, partly via volume transmission of ACh (Teles-Grilo Ruivo and Mellor, 2013). Increases in extracellular ACh are typically observed during exploratory behaviour in mice and drive activity in the theta to low-gamma range in CA1 and CA3 subfields through the activation of muscarinic ACh receptors (mAChRs) (Betterton et al., 2017; Fisahn et al., 1998). This is typically followed by an offline state characterised by sharp-wave ripples (SWRs) during immobility and slow-wave sleep, a time when ACh is low or absent (Zhang et al., 2021). Indeed, ACh has been shown to directly inhibit SWRs, allowing an animal to switch between a high-ACh, online attentive processing state dominated by low-gamma oscillations and a low-ACh, offline memory consolidation state dominated by SWRs (Jarzebowski et al., 2021; Zhang et al., 2021). This switch in the dynamics of hippocampal function has been proposed as a two-stage process of memory trace formation (Buzsaki, 1989). The cells responsible for these two network states have been suggested to map onto the two functional cell types described recently in CA3 (Hunt et al., 2018). Specifically, the highly adapting cells were proposed to initiate SWRs in low ACh conditions, whereas the non-adapting cells were proposed to be preferentially driven by a high ACh tone. However, whether cell types can switch behaviour to follow changes in cholinergic tone, therefore contributing to both network states, remains unexplored.

We have used organotypic hippocampal slices to study how chronic activation of cholinergic receptors modified the intrinsic properties of CA3 pyramidal neurons. We found that a highly plastic subpopulation of CA3 pyramidal neurons changed its intrinsic properties following long-term activation of AChRs. By increasing an M-type potassium current, chronic AChR activation transformed these cells from regular firing to hyperadapting neurons that typically fired a single, early action potential (AP) in response to current injection. This plasticity required neuronal activity and could be mimicked by specifically activating mAChRs. We provide evidence that through the modulation of an M-type potassium current, the numbers of CA3 neurons belonging to distinct functional classes can be altered in an activity-dependent manner. Our findings therefore suggest that the proportions of functionally-defined cell-types in the hippocampus is plastic.

## Results

We set out to assess the intrinsic forms of plasticity that take place following chronic stimulation of cholinergic pathways in CA3 hippocampal neurons. Using organotypic hippocampal slices we were able to emulate volume transmission of acetylcholine by applying the generic cholinergic agonist carbachol (CCH; 20 μM) extracellularly for 48 hours (Fig. 1A). Simultaneous extracellular recordings from CA1 and CA3 subfields showed that acute application of CCH induced a rapid increase in overall network activity and a shift in firing frequency towards the theta range. Inspection of extracellular recordings showed activity transitioned from sparse firing to high frequency bursts upon addition of CCH, in agreement with previous findings (Fig. S1A-B). The initial strong increase in activity decreased over time, as did the frequency bias, although neither returned to control levels within 48 hrs (Fig. S1C). This effect was not due to degradation or loss of CCH activity in the extracellular compartment, since acute application of the extracellular medium taken from cells incubated for 2 days in CCH showed similar increases in network activity (Fig. S1D). Together, our findings suggest that CCH treatment is able to produce robust increases in network activity that shift the firing properties of neurons into the theta frequency domain and that, although compensatory forms of plasticity are taking place in the network, they are not sufficient to completely overcome the strong CCH drive (Fig. S1E).

**Figure 1:**
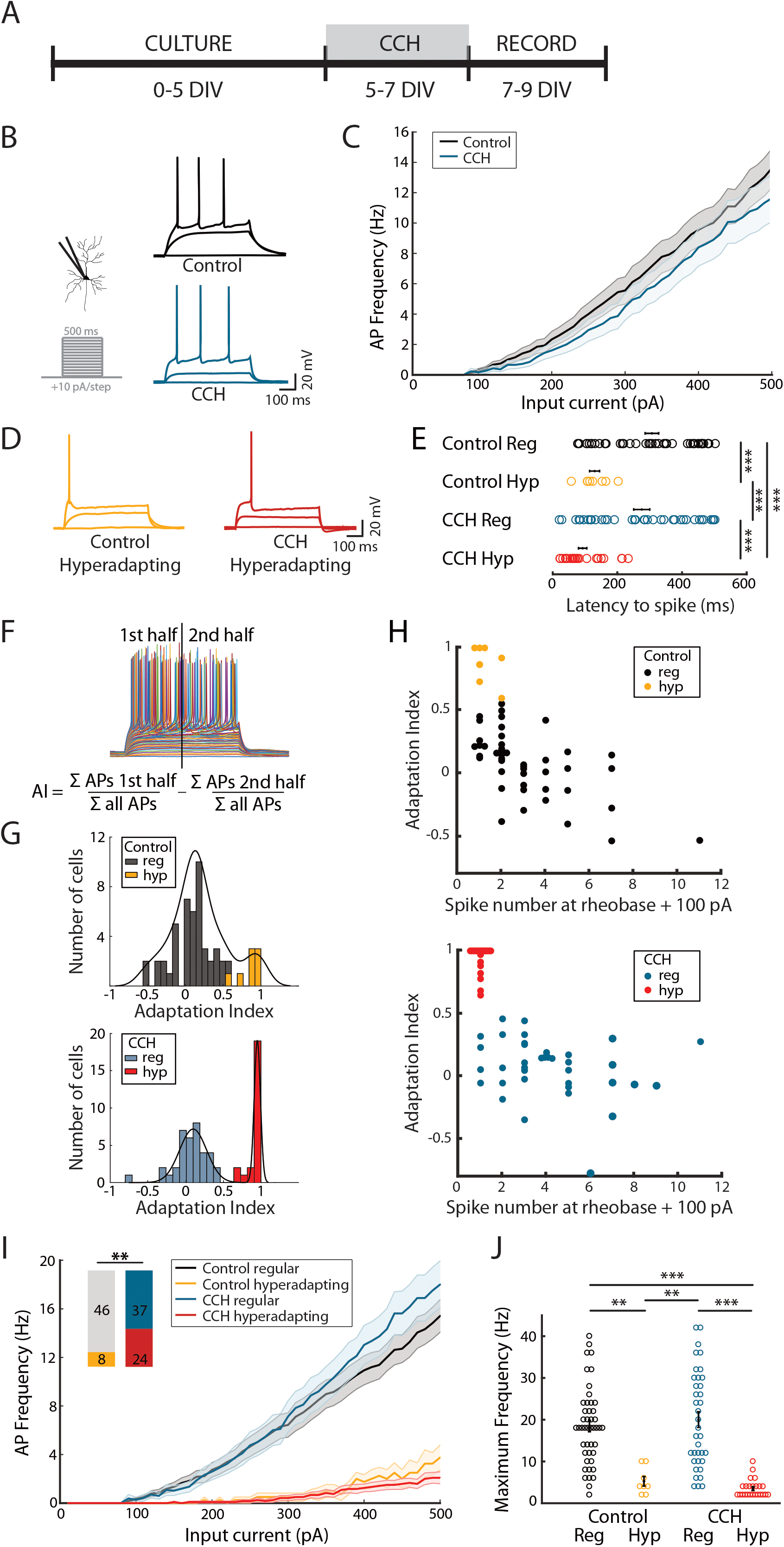
Chronic, cholinergic stimulation uncovers two distinct populations of CA3 pyramidal cells. **(A)** Timeline of preparation and treatment of organotypic hippocampal cultures. **(B)** Current injection protocol and two example voltage traces from regular spiking, control and carbachol (CCH) treated pyramidal cells. **(C)** Input-output curve for all control (black) and carbachol treated (blue) pyramidal cells. **(D)** Example voltage traces resulting from current injections in hyperadapting neurons in control and carbachol treated conditions. **(E)** Latency to first spike in control and carbachol treated neurons. *** p<0.001, one-way ANOVA followed by Tukey-Kramer test. **(F)** Calculation of adaptation index (AI) as the difference between the relative number of spikes in the first 250 ms of current injections and the second 250 ms of current injections. **(G)** Histogram of adaptation indices in control (top) and carbachol treated neurons (bottom), colour coded to indicate regular spiking and hyperadapting cells. **(H)** Scatter plot of adaptation index vs spike number at rheobase + 100 pA for control (top) and carbachol treated (bottom) pyramidal cells. Hyperadapting neurons with an adaptation index >= 0.6 and number of spikes at rheobase +100 pA < 4 are coloured yellow (top, control) and red (bottom, carbachol). **(I)** Input-output curves as in (c), split into hyperadapting and regular spiking neurons. Inset bars indicate the number of cells classed as hyperadapting and regular spiking in the control and treated conditions. ** p<0.01, Fisher’s exact test. **(J)** Maximum spike frequency in control and carbachol treated neurons. Error bars depict mean +/- S.E.M. ** p<0.01, *** p<0.001, one-way ANOVA followed by Tukey-Kramer test.

### Plasticity in the intrinsic excitability of a subset of CA3 pyramidal neurons by chronic ACh receptor activation

To assess the intrinsic excitability of CA3 neurons following chronic cholinergic stimulation, we performed whole-cell recordings in current-clamp mode in the absence of CCH. Current injection at the soma produced robust action potentials when rheobase was reached with neurons typically firing at higher frequencies with increasing current (Fig. 1B). The input-output curves showed that CCH-treated neurons, when pooled together, showed no obvious difference to untreated neurons (Fig. 1C) and only a small effect on rheobase (Fig. S2A). However, a closer look at the behaviour of individual CCH-treated neurons showed a large proportion of cells that fired only one, or very few, action potentials to currents that normally elicited repetitive spiking in untreated neurons (Fig. 1D). In addition, the APs occurred almost exclusively at the start of a current step (Fig. 1E), indicating strong adaptation. We termed these single-spiking cells hyperadapting neurons. The high proportion of hyperadapting neurons in CCH-treated slices prompted us to check whether these neurons were also present in control slices. We found a much smaller, but clearly identifiable, group of neurons that displayed a similar hyperadapting behaviour in untreated slices. We therefore divided neurons into either hyperadapting or regular spiking (Fig. 1F-I). To capture these firing features, we calculated an adaptation index where the fraction of spikes measured in the second half of a current step was subtracted from the first half, giving an overall measure of the temporal firing bias (Fig. 1F). A value of 1 indicates that all APs took place within the first half of a current step. Histograms of the adaptation index across all CCH-treated CA3 neurons uncovered two well-defined functional populations of cells (Fig. 1G), one of which showed high levels of adaptation that mirrored our intuitive observations (red bars in Fig. 1G). Furthermore, plots of adaptation index as a function of firing frequency (the latter taken at 100 pA beyond rheobase) showed that the cluster of highly adapting neurons typically fired a single AP (Fig. 1H) and only very high currents were capable of inducing multiple APs (Fig. S3). As expected, plots of the input-output functions for each group (Fig. 1I) showed that hyperadapting neurons had a higher rheobase (Fig. S2B) and a very low firing frequency compared to regular spiking neurons (Fig. 1J). We found that the firing properties of either hyperadapting or regular firing neurons were similar for control and CCH-treated slices (Fig. 1I), but the proportions of hyperadapting neurons increased substantially following CCH-treatment (from 15% to 40%; Fig. 1G-H). Intriguingly, a coarse map of the location of hyperadapting neurons in CA3 suggests that they are preferentially found in areas CA3a and CA3b, the two areas within the CA3 subfield that show the greatest amount of recurrent connections. Our data shows that there is a group of highly plastic CA3 neurons that is unmasked following CCH-treatment, resulting in an increase in hyperadapting cells.

We next set out to better understand the type of stimulus that led to the increase in the number of hyperadapting CA3 neurons. To establish whether activity (AP firing) is needed, we co-incubated slices with CCH and tetrodotoxin (TTX) to block all APs in the network during the CCH treatment (Fig. 2A). Current-clamp recordings found very few hyperadapting neurons following two days of incubation, suggesting this form of plasticity was dependent on neuronal/network activity (Fig. 2A-B). Since activity was clearly needed, we next decided to increase network activity via a pathway that would not involve cholinergic activation. Treatment with cyclothiazide (CTZ), an AMPA receptor modulator that prevents AMPA receptor desensitisation, induced large-scale increases in network activity. However, the pattern of network activity was markedly different from CCH treatment, with fewer bursts and a bias towards lower frequency regimes (not shown). Whole-cell recordings showed a rightward shift in the input-output curve after 48 hours of treatment with CTZ, as expected for a homeostatic decrease in excitability (Fig. 2C). However, we found no change in the proportion of hyperadapting neurons following this treatment (Fig. 2C-D). Together, our findings suggest that although activity is important, the emergence of hyperadapting neurons requires the activation of a cholinergic pathway.

**Figure 2:**
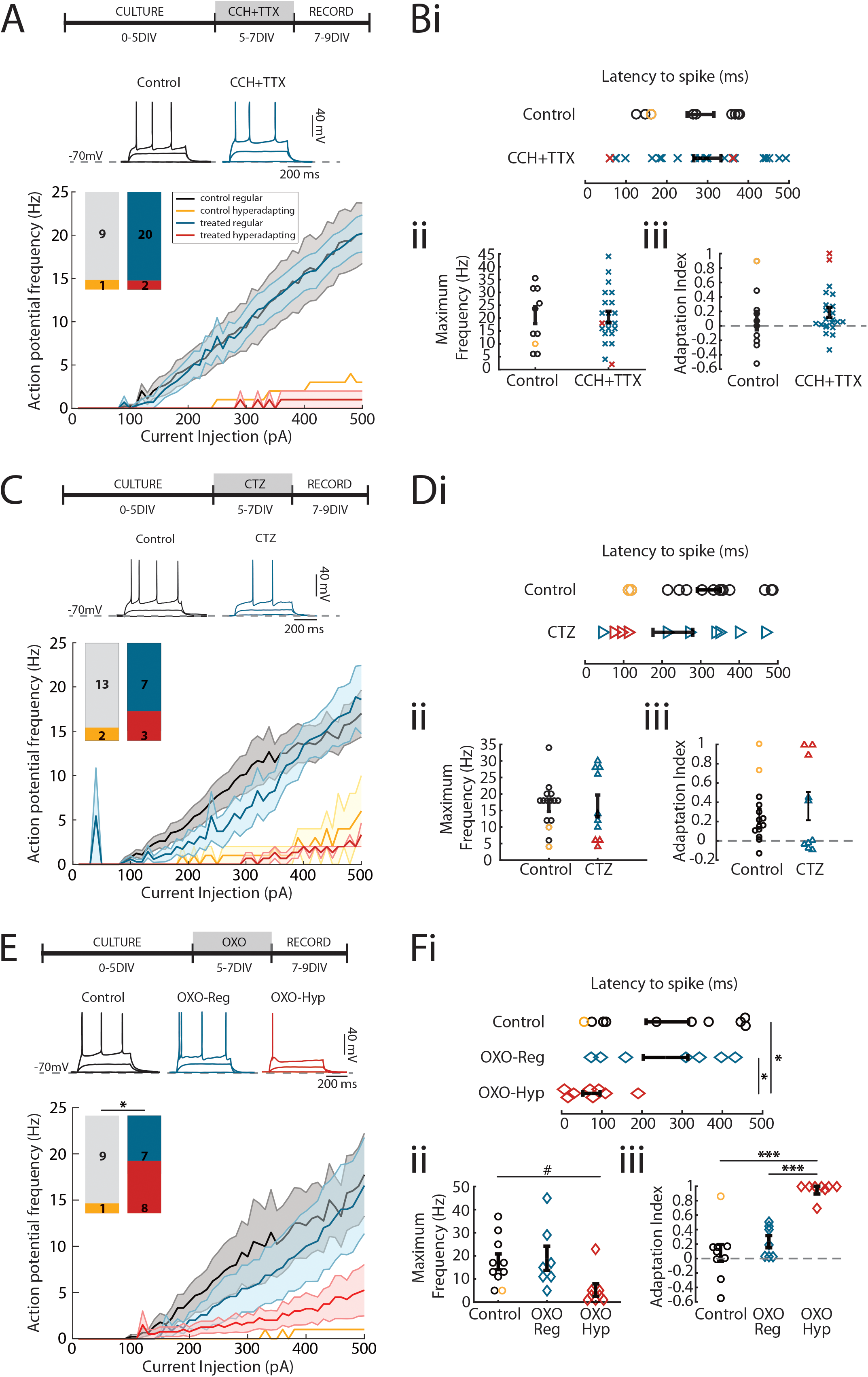
Plasticity of intrinsic excitability is activity dependent and triggered by muscarinic AChR activation. **(A)** Timeline of treatment with CCH (20 μM) and TTX (1 μM, top), example voltage responses to current injections (middle) and input-output curves split into hyperadapting and regular spiking neurons in control and CCH+TTX conditions (bottom). Bar graphs depict the number of cells in each category. **(Bi)** Latency to spike, **(Bii)** maximum action potential frequency and **(Biii)** adaptation index showing individual values and mean +/- S.E.M. **(C)** Timeline of treatment with cyclothiazide (5 μM, top), example voltage responses to current injections (middle) and input-output curves split into hyperadapting and regular spiking neurons in control and cyclothiazide conditions (bottom). Bar graphs depict the number of cells in each category. **(Di)** Latency to spike, **(Dii)** maximum action potential frequency and **(Diii)** adaptation index showing individual values and mean +/- S.E.M. **(E)** Timeline of treatment with oxotremorine-M (20 μM, top), example voltage responses to current injections (middle) and input-output curves split into hyperadapting and regular spiking neurons in control and oxotremorine-M conditions (bottom). Bar graphs depict the number of cells in each category. * p<0.05, Fisher’s exact test. **(Fi)** Latency to spike, **(Fii)** maximum action potential frequency and **(Fiii)** adaptation index showing individual values and mean +/- S.E.M. * p<0.05, *** p<0.001, oneway ANOVA followed by Tukey-Kramer test. # p<0.05, one-way ANOVA.

CCH is a broad cholinergic agonist that does not distinguish between nicotinic and muscarinic receptors. Whereas nicotinic AChRs drive increases in activity through ionotropic receptors, the muscarinic AChRs increase activity through intracellular second messenger cascades, which have been implicated in long-term forms of plasticity. We therefore tested whether activating mAChRs was sufficient to reproduce the phenotypes observed for CCH treatment (Fig. 2E). Indeed, we found that incubation with oxotremorine-M (OXO-M), a specific muscarinic acetylcholine receptor agonist, also uncovered a high proportion of hyperadapting neurons, similar to the numbers observed for CCH treatment (Fig. 2E-F). We concluded that activation of muscarinic ACh receptors is required for the appearance of hyperadapting CA3 neurons.

### Hyperadapting neurons do not show distinct morphological features

The fact that only a subset of neurons appears capable of becoming hyperadapting prompted us to look at whether there were any systematic differences in cell morphology between cells with different physiology. During recordings of neuronal output, some neurons were filled with Alexa-594 included in the patch pipette and imaged on a 2-photon microscope (Fig. 3A). 3D reconstructions of filled neurons did not show dramatic differences in dendritic morphology (Fig. 3B-D). We found that hyperadapting and repetitive firing cells had a similar dendritic length and arborisation index, as well as no significant biases towards basal or apical dendrites. A Sholl analysis also showed that there was no clear difference in dendritic branching nor any differences in the number of branchpoints (Fig. S4). Previous findings have reported a population of highly adapting burst-firing CA3 neurons that lack thorny excrescences (Hunt et al., 2018). We were unable to find clearly defined thorny excrescences in any of our neurons, which is likely due to the developmental stage of the circuit studied here (Ribak et al., 1985). Our results suggest that, at this stage in development, there appear to be no defining morphological features that would allow us to distinguish between the different functionally-defined neuronal cell types.

**Figure 3:**
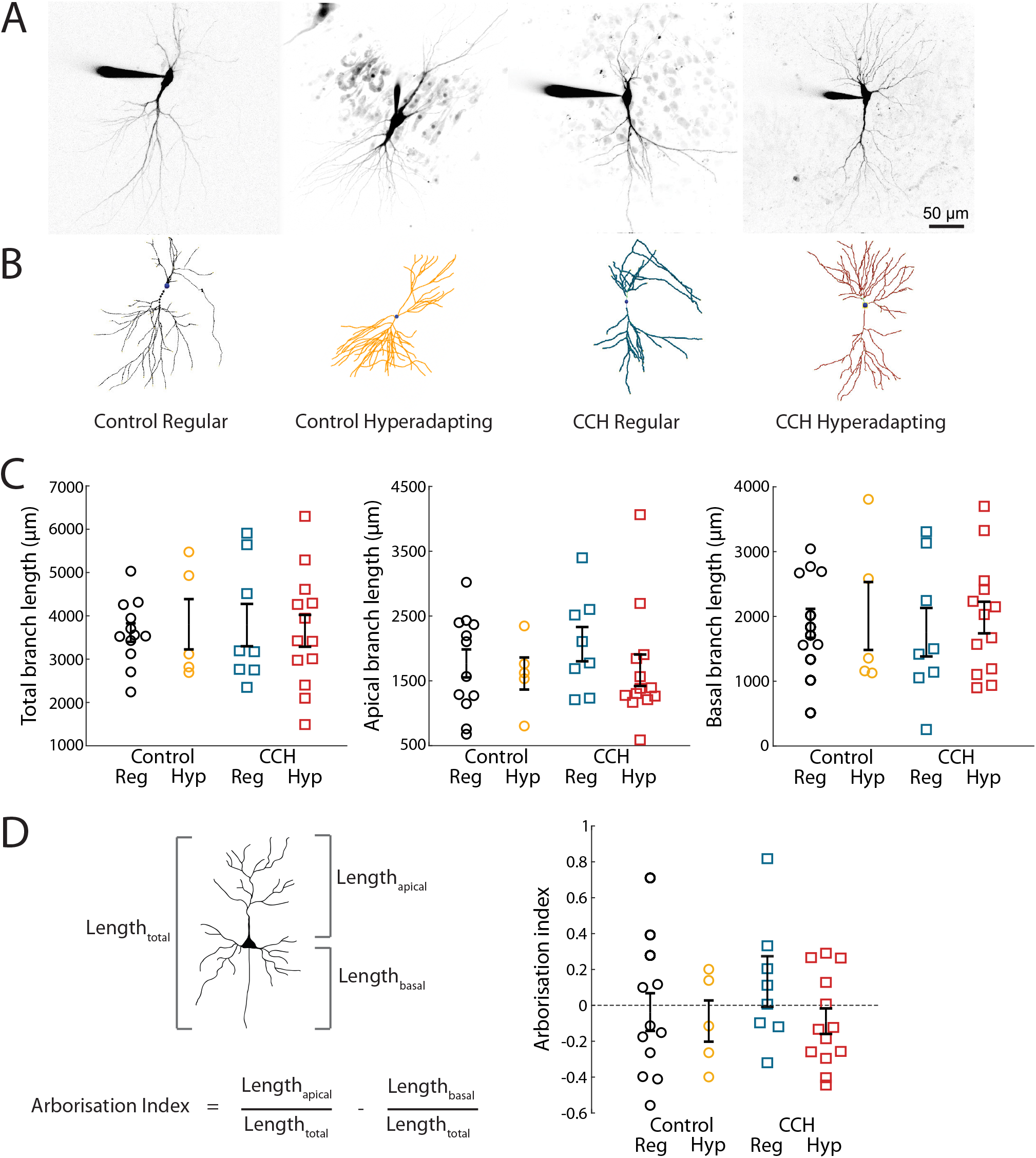
Hyperadapting and regular spiking CA3 neurons share the same morphological features. **(A)** Example images and **(B)** reconstructions of pyramidal cells filled with Alexa 594 through a patch pipette in control and carbachol-treated conditions. **(C)** Cumulative, dendritic branch length of the entire dendritic tree (left), the apical part of the dendritic tree (middle) and the basal part of the dendritic tree (right) **(D)** Calculation of arborisation index (left) and arborisation indices (right) in control and carbachol treated conditions. Error bars are mean +/- S.E.M.

### Upregulation of Kv7 activity generates hyperadapting neurons

In the hippocampus, activation of mAChRs have been shown to inhibit a slow, low voltage-activating, non-inactivating M-type K+ current (I_M_) carried by Kv7 channels (Brown and Adams, 1980; Selyanko et al., 2000). Since these channels have been implicated in adaptation during a spike train in a number of neurons, we speculated that they may also play a role in hyperadapting cells. We therefore performed voltage-clamp experiments to directly measure potassium currents in neurons previously identified in current-clamp mode as either hyperadapting or regular spiking (Fig. 4A-C). Since we needed to first establish the firing mode of each neuron, our macroscopic current measurements included all active conductances (Fig. 4A). To avoid contamination by the rapidly inactivating inward currents (carried mostly by Na^+^ and Ca^2+^ channels), we confined our measurements to the last 50 ms of the current trace, where I_M_ conductances are at their peak. We found that whereas the Kv7 antagonists XE-991 and Linopirdine had little effect on the steady-state outward K^+^ currents elicited in regular spiking neurons, hyperadapting neurons were highly sensitive, suggesting high levels of I_M_ in these neurons (Fig. 4B-C).

**Figure 4:**
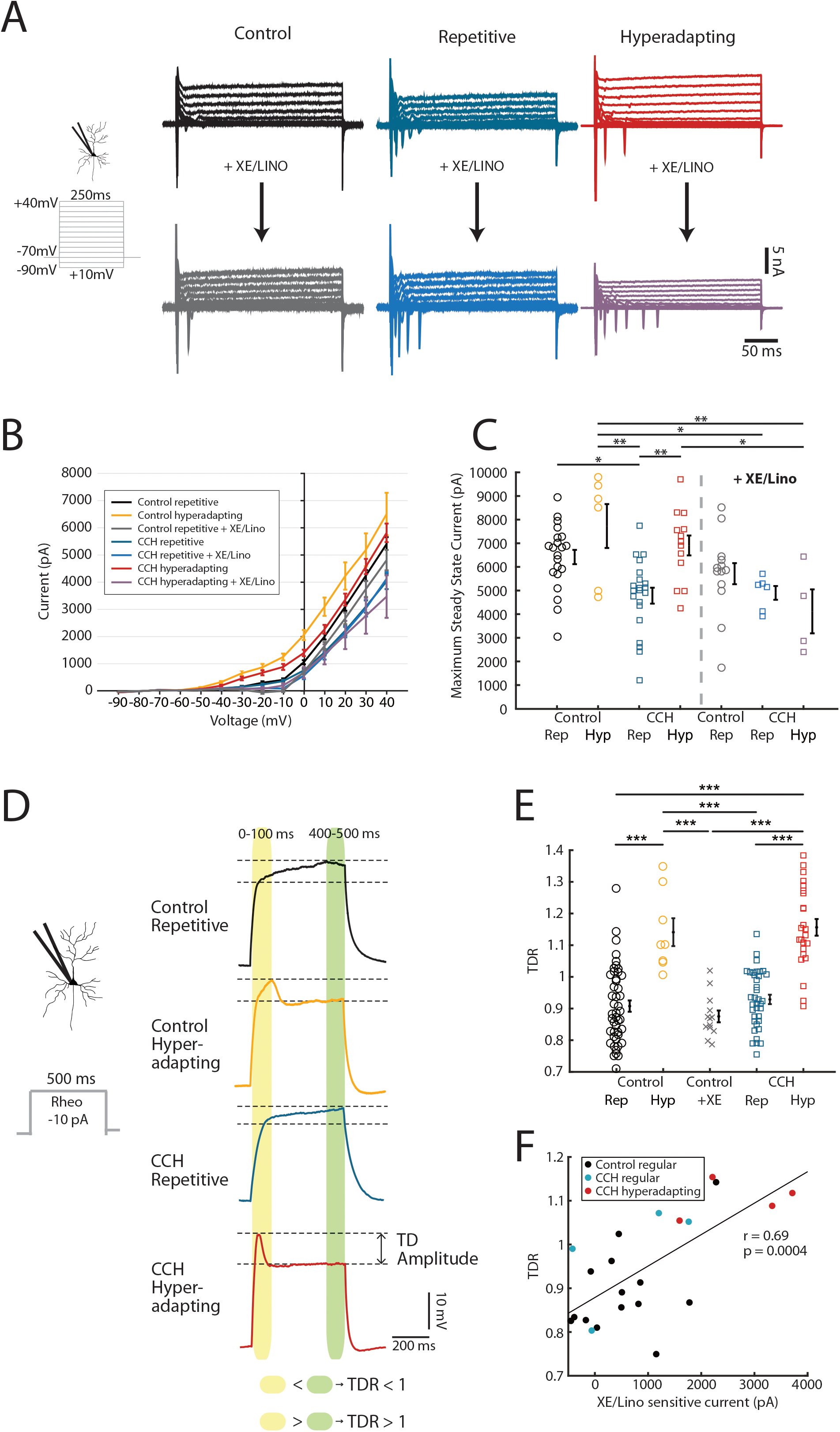
Hyperadapting neurons are characterised by an increased M-current. **(A)** Example voltageclamp recordings of control regular spiking (black), carbachol-treated, regular spiking (blue) and hyperadapting (red) neurons, before and after application of XE-991 and Linopirdine (20 μM each). Note the large steady-state current amplitude in hyperadapting neurons and the strong decrease with application of M-current blockers. **(B)** Current-voltage relationship for steady-state currents from the last 50 ms of each voltage step. **(C)** Maximum steady-state values at +40 mV across the different conditions and subtypes of neurons, with and without application of XE-991 and Linopirdine. Error bars are mean +/- S.E.M, * p<0.05, ** p<0.01. **(D)** Example voltage responses to a depolarising current pulse of 500 ms duration at 10 pA below rheobase, in the different conditions and subtypes of neurons. The transient depolarisation ratio (TDR) was calculated as the ratio of the mean amplitude in the first 100 ms and the mean amplitude in the last 100 ms. **(E)** Transient depolarisation ratio showing individual values and mean +/- S.E.M. *** p<0.001, one-way ANOVA followed by Tukey-Kramer test. **(F)** Scatter plot of TDR vs XE-991/Linopirdine sensitive current with least squares fit line, Pearson’s correlation coefficient and p-value. XE/Lino sensitive current was calculated as the steady state current at +40 mV after XE-991/Linopirdine application subtracted from steady state current at +40 mV before application of channel blockers.

To further investigate this, we switched to examining the voltage traces measured under current-clamp recordings (Fig. 4D). Hyperadapting neurons in both control and CCH-treated slices regularly showed a clearly identifiable transient depolarisation (TD) in the initial part of the voltage trace at subthreshold potentials, measured as transient depolarisation ratio (TDR; Fig. 4D), which was not present in regular firing neurons (Fig. 4D-E). The low voltage activation profile of the I_M_ conductance, combined with its slow onset kinetics, suggested that this transient depolarisation may be a signature for the presence of I_M_. Indeed, we found that this transient depolarisation correlated well with the I_M_ currents measured above (Fig. 4F). Together, our data strongly suggests that the mechanism behind the appearance of hyperadapting neurons is driven by an increase in an I_M_ conductance, which exerts a strong effect on the shape of the voltage response and explains its high levels of adaptation.

To specifically test for this, we treated hyperadapting neurons from CCH-treated slices with XE-991 and Linopirdine acutely (Fig. 5A). In line with our findings, the single AP firing profile switched to a regular firing mode immediately upon addition of the antagonists and resulted in a normalisation of the input-output function (Fig. 5B). In direct agreement with our findings above, both the adaptation index and the TDR were strongly reduced and, as expected from blocking a K^+^ channel that is open at rest, we also observed an increase in both the resting membrane potential and input resistance (Fig. 5C). Since I_M_ is typically inhibited by acetylcholine acting on mAChRs (Brown and Adams, 1980), we treated hyperadapting cells acutely with CCH at the same concentration as used in the chronic treatment. Once again, we found that hyperadapting neurons converted to a regular spiking behaviour upon addition of CCH (Fig. 5D-E), which we found also dramatically reduced the adaptation index and TDR (Fig. 5F). We conclude that a subset of CA3 neurons strongly increase an I_M_ conductance in response to chronic activation of mAChRs, which switches their output to a hyperadapting modality once the cholinergic tone is removed. The unmasking of hyperadapting neurons after the removal of ACh suggests that this plasticity is unlikely to play a homeostatic role in the recovery of network activity while ACh is still present. Instead, this finding points to an instructive role in the emergence of hyperadapting neurons, where exposure to cholinergic activity converts all neurons into regular firing cells but then leaves behind a large subset of neurons with very different physiological properties once ACh subsides. It also shows that hyperadapting neurons are highly sensitive to cholinergic activity and can readily switch firing modalities acutely – from regular firing in the presence of acetylcholine, to hyperadapting in its absence.

**Figure 5:**
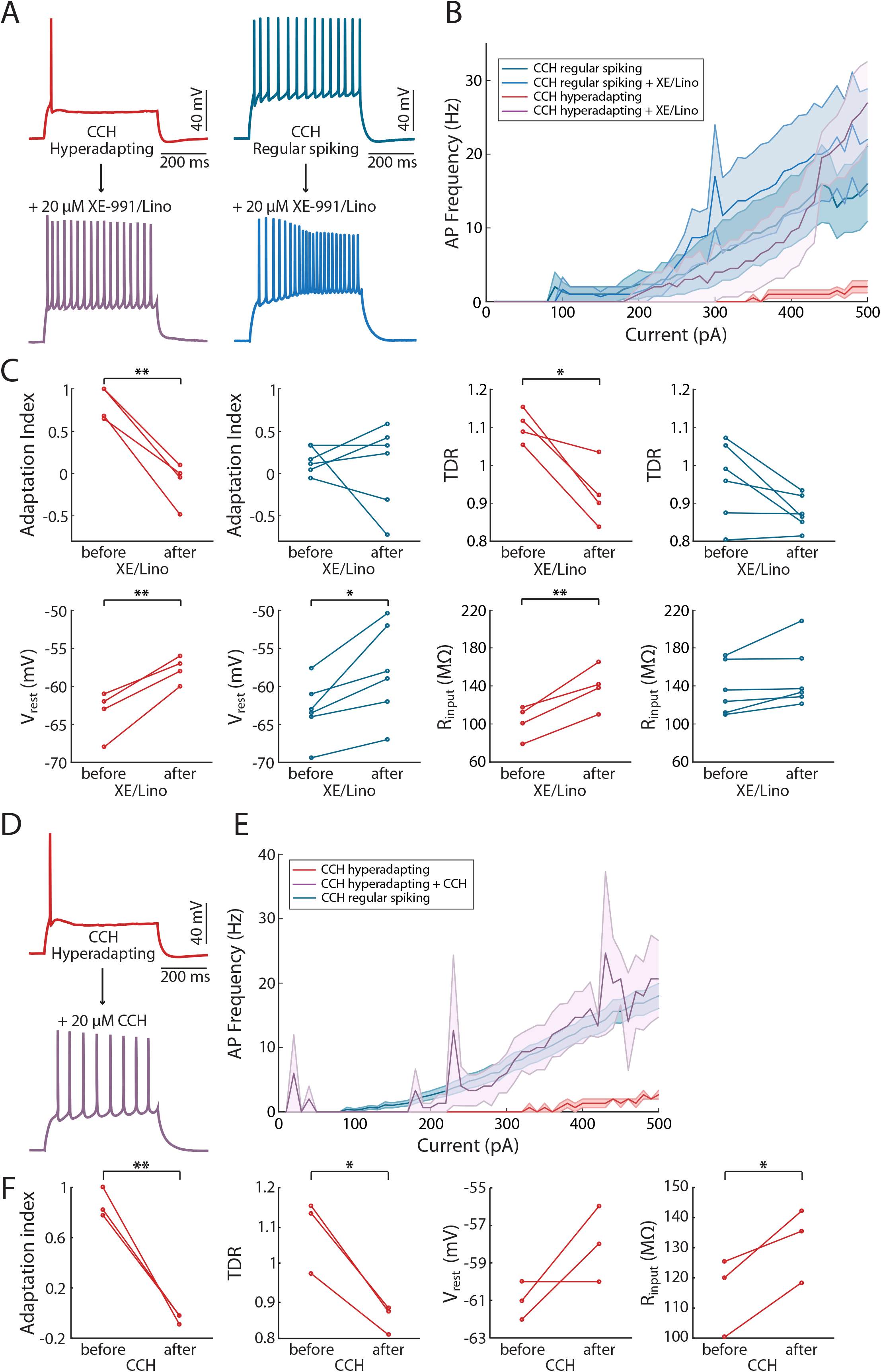
Acute cholinergic activation converts hyperadapting into regular spiking neurons. **(A)** Example recordings of action potentials in response to 500 ms current injections, in carbachol-treated hyperadapting and regular spiking neurons, before and after application of XE-991/Linopirdine. **(B)** Input-output curves showing action potential frequency in response to increasing current injections in carbachol-treated, hyperadapting and regular spiking neurons, before and after application of XE-991/Linopirdine. **(C)** Plots showing the changes in adaptation index, transient depolarisation ratio, resting membrane potential and input resistance with application of XE-991/Linopirdine. Error bars depict mean +/- S.E.M, * p<0.05, ** p<0.01, paired-sample Student’s t-test. **(D)** Example recordings of action potentials in response to 500 ms current injections, in a carbachol-treated hyperadapting neuron, before and after acute application of carbachol (20 μM). **(E)** Input-output curves showing action potential frequency in response to increasing current injections in carbachol-treated neurons before and after acute application of carbachol. **(F)** Plots showing the changes in adaptation index, transient depolarisation ratio, resting membrane potential and input resistance with acute application of carbachol. Error bars depict mean +/- S.E.M, * p<0.05, ** p<0.01, paired-sample Student’s t-test.

## Discussion

We show that long-term exposure to cholinergic agonists increases the number of CA3 neurons that show a strong adaptation in firing frequency, typically eliciting a single AP at the onset of a long current pulse. Whereas these hyperadapting neurons were rarely found in control slices (~15% of neurons patched), their numbers increased dramatically (~40% of neurons patched) following 48hrs treatment with cholinergic agonists. The appearance of hyperadapting neurons required spiking activity and the specific activation of mAChRs, resulting in the upregulation of an M-type potassium current. We propose that the neuromodulator acetylcholine can tweak the response properties of a subset of CA3 neurons, altering the relative numbers of functionally defined neuronal cell types in the hippocampus.

### Neuronal subtypes in CA3

Recent work has shown that there are at least two different types of CA3 pyramidal neurons in the hippocampus with distinct functional properties. Although the majority of CA3 neurons showed regular firing patterns that increased in frequency with increasing current, a subset of pyramidal cells displayed more transient firing responses with high frequency bursts at the onset of a pulse, that rapidly adapted (Hunt et al., 2018). The latter, referred to as ‘bursting’, represented roughly 30% of CA3 pyramidal cells and were predominantly found in distal areas of CA3, including CA3A and CA3B. Functionally, these bursting neurons have very similar characteristics to the hyperadapting neurons described here. Both showed strong adaptation, are found in similar areas of the CA3 subfield and accounted for a similar fraction of the total pyramidal cell population. However, since hyperadapting neurons did not fire bursts, but typically a single AP, we were reticent to call them bursting, even though they likely represent an extreme form of the adaptation seen in bursting neurons. Morphologically, bursting neurons showed a unique feature that differentiates them from regular spiking neurons – the absence of thorny excrescences (Hunt et al., 2018). In our young organotypic slices (cultured at P7 and grown for ~1 week), we could not detect any clearly defined thorny excrescences, which may reflect the fact that these synaptic protrusions develop relatively late (Ribak et al., 1985). We also found no difference in dendritic morphology between hyperadapting and regular spiking neurons. Although only speculative, it is likely that the two populations of cells described here match those described previously (Hunt et al., 2018), but that our neurons are not sufficiently developed to show the distinct dendritic morphologies observed in more mature neurons in intact tissue.

### A mechanism for the emergence of hyperadapting neurons

Our results show that the upregulation of an M-type current is responsible for the distinctive firing properties of hyperadapting neurons. M-currents are produced by Kv7 channels (Brown and Passmore, 2009) and their modulation controls the output of CA3 pyramidal neurons (Brown and Randall, 2009; Hunt et al., 2018). Importantly, Kv7 activity is tightly modulated by mAChRs (Selyanko et al., 2000). In general, activation of mAChRs inhibits Kv7 channels through a G-protein cascade that leads to increased activity in the hippocampus and the appearance of theta/gamma oscillations (Teles-Grilo Ruivo and Mellor, 2013). Furthermore, Kv7 channel expression is highly plastic (Baculis et al., 2020). Acute induction of epileptic activity in hippocampal neurons increases the expression of Kv7 transcripts (KCNQ2 and 3) in what appears to be a homeostatic mechanism to normalise activity levels (Zhang and Shapiro, 2012). Conversely, chronic decreases in activity achieved by blocking NMDA receptors reduced Kv7 currents (and transcripts) in dissociated hippocampal neurons (Lee et al., 2015), showing that Kv7 channels appear to modulate their expression bidirectionally in a homeostatic manner. Other forms of plasticity have also been linked to Kv7 channel expression. For example, sustained stimulation of cholinergic inputs in hippocampal slices resulted in an increase in the intrinsic excitability of dentate granule cells, mediated by the long-term inhibition of a Kv7 current (Martinello et al., 2015). This form of plasticity required the activation of T-type calcium channels, suggesting the need for calcium influx.

We found that the increase in Kv7 activity in hyperadapting neurons was not simply caused by increases in overall levels of activity, but instead required the specific activation of mAChRs, together with increased spiking activity, for this plasticity to take place. However, our findings do not tally well with a homeostatic interpretation of Kv7 upregulation. The increase in network activity observed when AChR agonists were delivered did not show any obvious adaptation during the 2 days of cholinergic stimulation, despite the increase in Kv7 currents. This is likely due to the fact that Kv7 channels are blocked during the activation of mAChRs, so that increases in Kv7 conductances will only become apparent once cholinergic stimulation ceases. Indeed, hyperadapting neurons are only observed in the absence of ACh agonists and either activation of AChRs or acute block of Kv7 channels reverts them to a repetitive firing mode. As a result, this form of plasticity did not allow neurons to adapt to the stimulus, but instead appeared to prime them for when ACh levels decreased (Fig. 6). This differs from recent work in dissociated neuronal cultures where chronic inhibition of Kv7 channel activity with XE-991 or with Carbachol caused a homeostatic decrease in excitability that relied on the relocation of the AIS, which includes Kv7 channels, to dampen firing during drug application (Lezmy et al., 2017). We did not observe a substantial decrease in CA3 pyramidal cell excitability during chronic AChR activation in our experiments, suggesting that multiple forms of plasticity may be elicited, depending on cell type and/or neuronal preparation.

**Figure 6:**
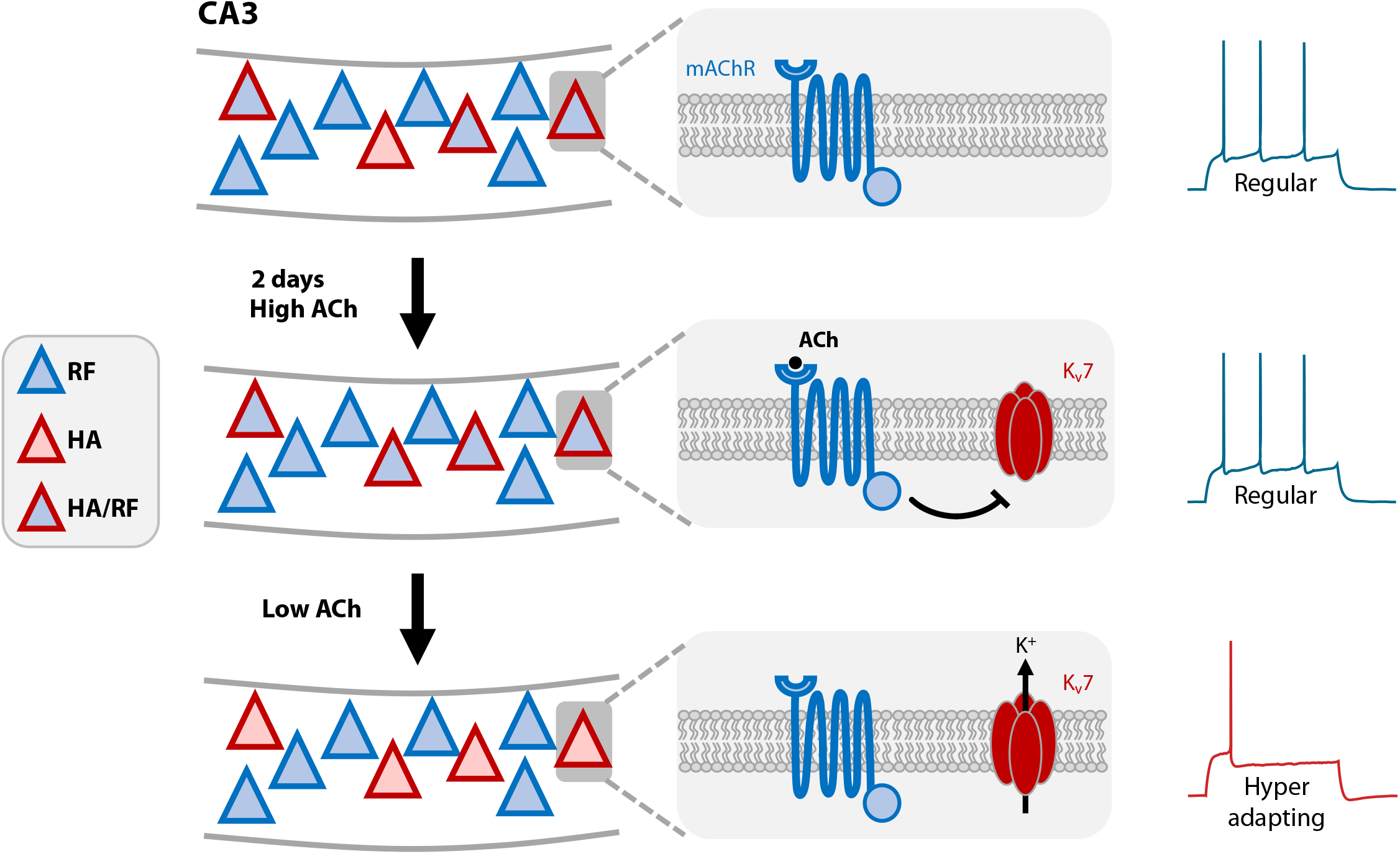
Possible mechanism describing the modulation of intrinsic excitability by cholinergic stimulation in a subset of CA3 pyramidal neurons. CA3 pyramidal cells consist of regular firing (RF, blue-filled triangles) and hyperadapting neurons (HA, red-filled triangles). A subset of neurons are able to switch firing behaviour between regular and hyperadapting, depending on cholinergic activity (HA/RF, triangle with red outline and blue fill). **Top**: In baseline conditions, when cholinergic activity is low, HA/RF neurons fire action potentials in a regular manner and express low amounts of Kv7 channels. **Middle**: During chronic cholinergic stimulation, HA/RF neurons increase their levels of Kv7 channel conductance, but remain regular firing whilst ACh remains. This is likely due to the direct block of Kv7 by mAChRs. **Bottom**: Once the intense cholinergic activity state has subsided, the drop in extracellular ACh relieves the block of Kv7 channels by mAChRs, causing these neurons to become hyperadapting.

### A possible role in brain function

The diversity of CA3 hippocampal neurons has been linked to the process of learning and memory consolidation (Hasselmo, 2006). Cholinergic inputs from the medial septum are thought to play a key role in this process (Teles-Grilo Ruivo and Mellor, 2013). Typically, ACh levels in the rodent hippocampus are high during exploratory behaviour and decrease substantially during immobility or slow-wave sleep (Betterton et al., 2017; Fisahn et al., 1998; Jarzebowski et al., 2021; Zhang et al., 2021). Both these phases are required for the acquisition and consolidation of learnt behaviours (Buzsaki, 1989). During high ACh tones, as the animal is exploring, hippocampal activity increases in the theta/low gamma range through the activation of muscarinic ACh receptors and SWRs are rarely observed. This contrasts with the offline states seen during inactivity, where ACh tone is low and SWRs are observed. Recent work has proposed that bursting cells contribute to initiating SWRs during the low ACh state, whereas increases in activity during the high ACh state is mediated by regular spiking cells (Hunt et al., 2018). The hyperadapting neurons described here provide an elegant solution to the two behavioural states described above. During high ACh tone, hyperadapting cells will behave like regular spiking cells, since their Kv7 channels will be inhibited. When ACh tone drops, their highly adapting firing properties will be unveiled allowing them to contribute to SWRs. This acute switch in firing properties of hyperadapting cells makes them good candidates for explaining the transitions in activity observed between different brain states in the hippocampus. The fact that the number of hyperadapting cells depends on the past history of cholinergic activity opens up the possibility that the types of activity for different brain states is plastic. A prolonged ACh tone, which would lead to more hyperadapting neurons, could therefore prime the network for altered network activity (e.g. presence of SWRs) in the low ACh (offline) state. Our findings show that ACh, a key neuromodulator in the brain, can alter the relative proportions of cell types in the CA3 in a use-dependent manner, with important implications for network dynamics.

## Acknowledgements

We would like to thank Matthew Grub, Guilherme Neves, Rachel Jackson and Mala Shah for useful feedback and comments on the manuscript. This research was funded in whole, or in part, by the Wellcome Trust (095589/Z/11/Z and 215508/Z/19/Z). For the purpose of open access, the author has applied a CC BY public copyright licence to any Author Accepted Manuscript version arising from this submission. This work was also supported by an ERC Starter Grant (282047) and a BBSRC project grant (BB/S000526/1) to JB, as well as an MRC studentship to CP.

## Methods

### Organotypic slice preparation

Organotypic hippocampal slice cultures were prepared as in (Stoppini et al., 1991). Hippocampi of P7 male Sprague-Dawley rats (Charles River, UK) were dissected in cold GBSS supplemented with D-Glucose (34.7 mM) and cut into 400-μm-thick slices using a McIlwain tissue chopper. Slices were placed onto Millicell-CM membranes and maintained in culture media composed of 25% (vol/vol) EBSS (Invitrogen), 49% (vol/vol) MEM (Invitrogen), 1% (vol/vol) B27 (Invitrogen), 25% (vol/vol) heat-inactivated horse serum (PAA), and 6.2 g/l glucose (Fisher). Slices were incubated at 36°C and 5% CO2. After 4-6 days in vitro, carbachol (20 μM), carbachol+TTX (20 μM + 1 μM), cyclothiazide (5 μM), or Oxotremorine-M (20 μM) was added to the slice media. Slices were then cultured in the drugs for a further 48 hours prior to recordings. Control slices were paired with treated slices to ensure similar age at recording.

### Electrophysiology

Slices were transferred to a recording chamber and continuously perfused with artificial cerebrospinal fluid (ACSF) at 30°C. ACSF contained 119 mM NaCl, 2.5 mM KCl, 1 mM MgCl2, 3 mM CaCl2, 1 mM NaH2PO4, 26.2 mM NaHCO3, and 11 mM D-Glucose. ACSF was constantly oxygenated using carbogen (5% CO2, 95% O2). Patch pipettes made from thick-walled borosilicate glass capillaries with an inner filament (1.5mm inner diameter, 0.86mm inner diameter; Sutter Instruments, Novato, CA, USA) were pulled on a P-97 Flaming/Brown Micropipette Puller (Sutter Instruments). All recordings were made with a Multiclamp 700B amplifier (Molecular Devices), Bessel filtered at 10 kHz, digitized and sampled at 50 kHz. All recordings were made using the pClamp software program. Neurons were visualised with a Scientifica two-photon microscope using either a 10x 0.25NA Olympus air objective or a 40X 0.8NA Olympus water immersion objective and Dodt Gradient Contrast.

### Extracellular recordings

For extracellular recordings pipettes were pulled to ~1MΩ and filled with ACSF. Pipettes were placed into the stratum pyramidale of CA3 and CA1. Recordings of voltage signals were made in the current clamp configuration with no current injected (I = 0). Recordings were made from slices in 30-minute segments up to a maximum of 2 hours, at which point slices were discarded. Recordings were notch filtered offline to remove 50 and 60 Hz noise.

### Whole-cell recordings and analysis

For whole cell recordings, 3-6 MΩ pipettes were pulled, fire-polished and filled with internal solution that contained: 120 mM K-Gluconate, 28.5 mM sucrose, 10 mM HEPES, 9 mM KCl, 10 mM KOH, 4mM Na2ATP, 0.4 Na2GTP. Internal solution was adjusted to pH 7.3 using KOH. For some experiments Alexa-594 (Thermo Fisher Scientific, MA, USA) was added to the internal to 20 μM. To block synaptic transmission, 25 μM AP-V, 20 μM SR95531 and 10 μM NBQX were added to the ACSF. For certain experiments, we also added 20 μM XE-991 and 20 μM Linopirdine or 20 μM CCH, as stated in the text. CA3 neurons were identified based on their position in the slice and CA3 was subdivided into subregions using the criteria set out in (Li et al., 1994). Confirmation that pyramidal neurons had been targeted was made during recordings by checking the magnitude of the sag potential and the maximum firing frequency during 500 ms current injections. CA3 neurons would be expected to have little to no sag current and a firing frequency of <50 Hz. Cells that had an initial V_rest_ > −55 mV or R_series_ > 30 MΩ were discarded. For longer recordings which involved changing drugs in the bath, any cells which had a change in R_series_ greater than 10% were discarded. Reported values are not corrected for liquid junction potential. The resting membrane potential was measured immediately after membrane breakthrough in current clamp in I = 0. For current clamp recordings, all recordings were bridge balanced and pipette capacitance was neutralised. A bias current was injected to hold the membrane voltage of each cell at −70 mV. 500 ms duration current steps of increasing magnitude (+10 pA) were injected via the patch pipette in order to measure the spiking properties. The transient depolarisation ratio (TDR) was calculated from the current step just prior to rheobase (Fig. 4D) as:

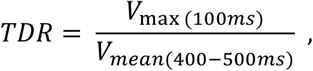

where V_max(100ms)_ is the maximum voltage recorded during the first 100 ms of the current injection and V_mean(400-500ms)_ is the mean voltage during the last 100 ms of the 500 ms current injection. Latency to spike was calculated at rheobase as the time it took from the beginning of the current injection to the first action potential. Adaptation index (AI) was calculated over all sweeps for a given cell which contained at least one spike and was defined as (see also Fig. 1F):

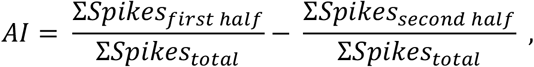

where ΣSpikes_first half_ is the sum of all action potentials that occurred in the first 250 ms of current injections and ΣSpikes_second half_ is the sum of all action potentials that occurred in the last 250 ms of current injections. For voltage clamp recordings, series resistance compensation was performed in order to achieve an effective R_series_ of 7 MΩ. R_series_ was monitored throughout experiments and adjusted if any changes occurred. If R_series_ increased by more than 10%, the recordings were discarded. Voltage steps of 250 ms duration were applied in 10 mV increments from −90 mV to +40 mV. Leak current was subtracted offline by calculating the passive leak current from −90 mV to −60 mV and fitting a curve to these values. Passive leak current for each voltage step was then calculated as the voltage step multiplied by the slope of the curve and subtracted from the recorded current value. Steady-state outward current was calculated as the mean current from the last 50 ms of each sweep.

### Imaging and image analysis

Neurons were filled with Alexa 594 through the patch pipette. Image stacks of Alexa 594-filled neurons were taken ~15 minutes after achieving whole-cell configuration, using a two-photon microscope (Scientifica) equipped with a 40x 0.8 NA water immersion objective and excited with a Chameleon femtosecond laser (Coherent) at 810 nm. Images were centred over the soma of the recorded cell and z-stacks were taken using Scanimage software. Neurons were traced using NeuTube software (Feng et al., 2015). SWC files generated from tracing were analysed using the TREES toolbox software package in MATLAB (Cuntz et al., 2010). Sholl analysis was carried out with concentric rings of increasing diameter in steps of 20 μm centred on the soma. All other morphological parameters were extracted using the TREES toolbox. For each cell the analysis was performed over the entire dendritic tree, as well as separately on the apical and basal arbours. Arborisation index was calculated for each traced neuron and defined as (see also Fig. 3D):

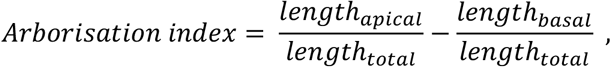

where length_apical_ is the total length of all apical dendrites, length_basal_ is the total length of all basal dendrites and length_total_ is the total length of the entire dendritic tree.

### Statistics

For all statistical tests, normality of the data was first determine by using Shapiro-Wilks test with a threshold of p=0.05. For normal distributions, t-tests were used; for non-normal distributions, Wilcoxon rank sum tests were used. In both instances the threshold for significance was p=0.05. For comparisons between more than two groups, one-way ANOVAs were used with post-hoc Tukey-Kramer tests.

