## Supplemental Figures for "Cholinergic stimulation modulates the functional composition of CA3 cell types in the hippocampus"

### Supplemental Figure Legends

**Supplemental Figure 1: Hippocampal population activity increases mainly in the low-gamma band with carbachol application. (A)** Spontaneous population activity recorded extracellularly, simultaneously in the CA1 (top) and CA3 (bottom) subregions of untreated hippocampal organotypic slices. Dotted region marked in the left recording traces depict the part that is enlarged in the middle. Right: Spectrograms for the recordings depicted on the left. **(B)** Extracellular recording of spontaneous population activity as in (A), in untreated slices after acute application of carbachol (20 μM). **(C)** Extracellular recording of spontaneous population activity as in (A), in slices treated with carbachol for 48 hours prior to recording. **(D)** Extracellular recording of spontaneous population activity as in (A), in untreated slices after acute application of carbachol that was incubated for 48 hours in growth medium of a different set of organotypic slices. **(E)** Power spectra of the CA1 and CA3 recordings shown in (A-D).

**Supplemental Figure 2: Action potential threshold is higher in hyperadapting than regular spiking pyramidal neurons. (A)** Rheobase calculated from 500 ms current injections in control and carbachol-treated pyramidal neurons. Error bars depict mean +/- S.E.M., ***$p<0.001$, Student's t-test. **(B)** Rheobase measurements as in (A), but split by hyperadapting and regular spiking phenotype. Error bars depict mean +/- S.E.M., * $p<0.05$, ***$p<0.001$, one-way ANOVA followed by Tukey-Kramer test.

**Supplemental Figure 3: Input-output curves show a clear separation between hyperadapting and regular spiking neurons, even at high levels of current injections.** Input-output curves showing action potential frequency in response to increasing current injections for each individual neuron, in control (left) and carbachol-treated (right) conditions, colour-coded according to functional cell type.

**Supplemental Figure 4: Morphological features of CA3 pyramidal cell subpopulations. (A)** Number of branchpoints in the entire dendritic tree (left), and split into apical dendritic tree (middle) and basal dendritic tree (right). **(B)** Mean branch order of the entire dendritic tree (left), and split into apical dendritic tree (middle) and basal dendritic tree (right). **(C)** Sholl analysis of the entire dendritic tree (left), and split into apical dendritic tree (middle) and basal dendritic tree (right).

**Figure S1**

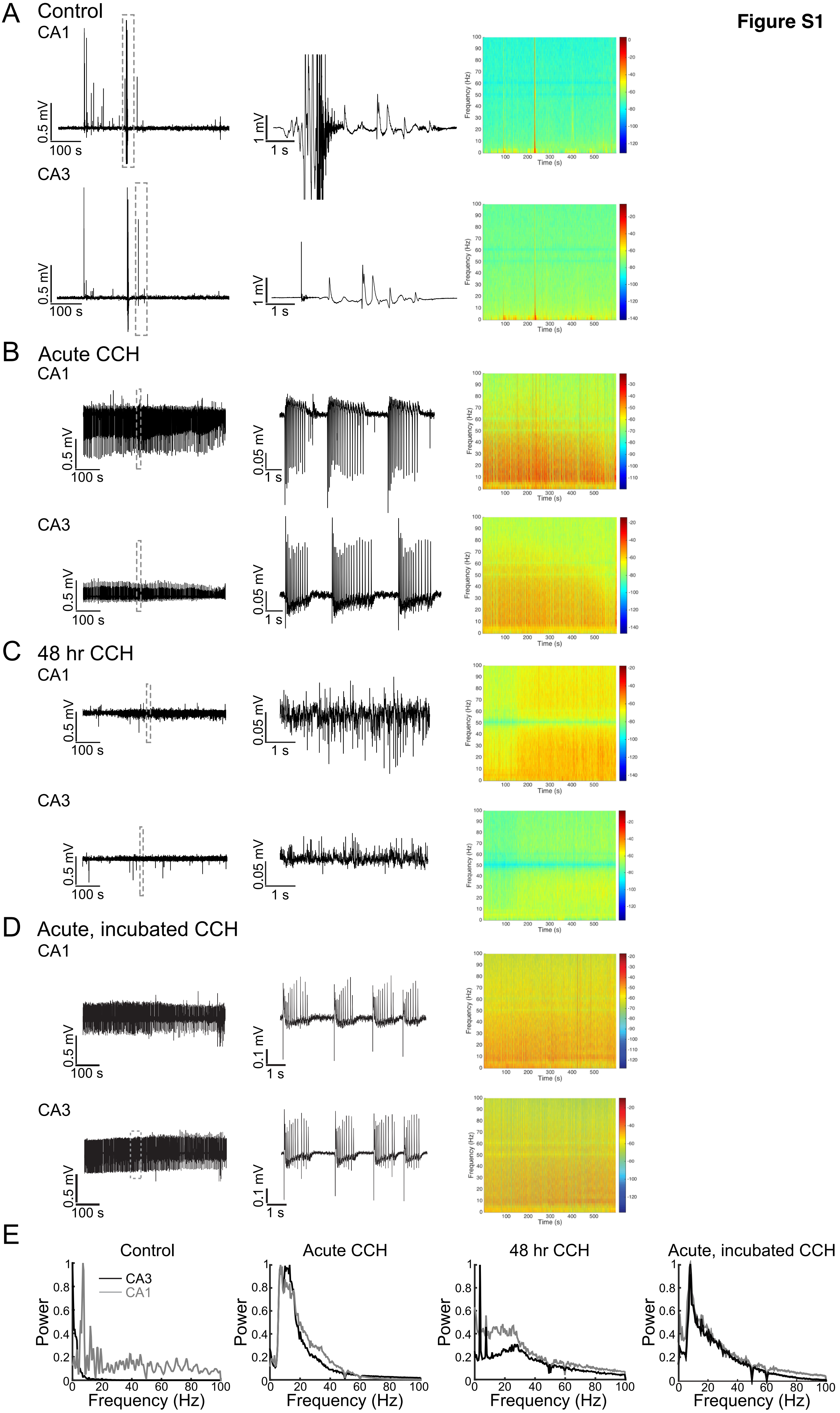
A
Control
CA1
CA3
0.5 mV
100 s
1 mV
1 s
Frequency (Hz)
Time (s)
B
Acute CCH
CA1
CA3
0.05 mV
C
48 hr CCH
D
Acute, incubated CCH
0.1 mV
E
Control
Acute CCH
48 hr CCH
Acute, incubated CCH
CA3
CA1
Power
Frequency (Hz)

**Figure S2**

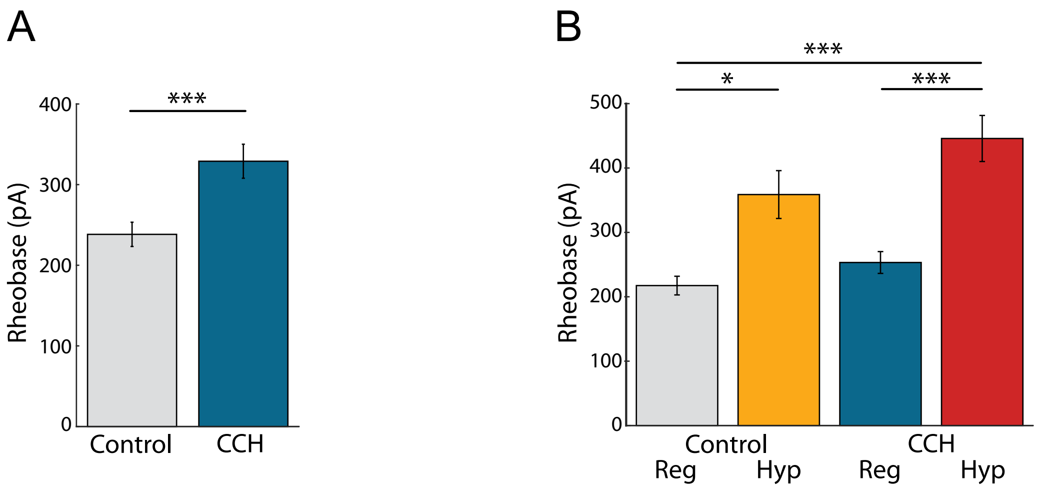
A
***
400
300
200
100
0
Rheobase (pA)
Control
CCH
B
***
*
***
500
400
300
200
100
0
Rheobase (pA)
Control
CCH
Reg
Hyp
Reg
Hyp

**Figure S3**

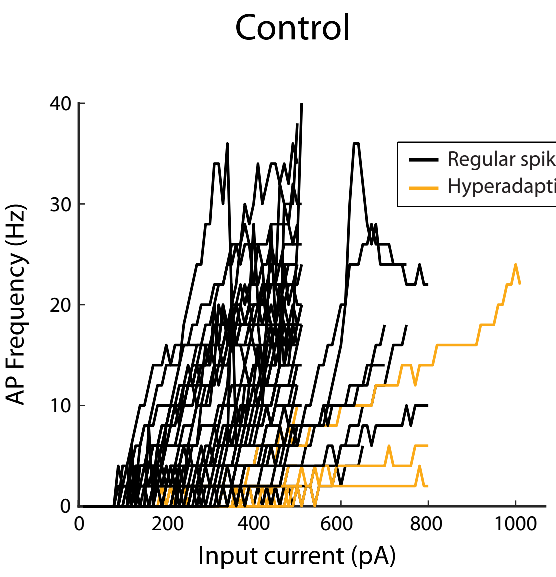
Control
40
30
20
10
0
AP Frequency (Hz)
0
200
400
600
800
1000
Input current (pA)
Regular spiking
Hyperadapting

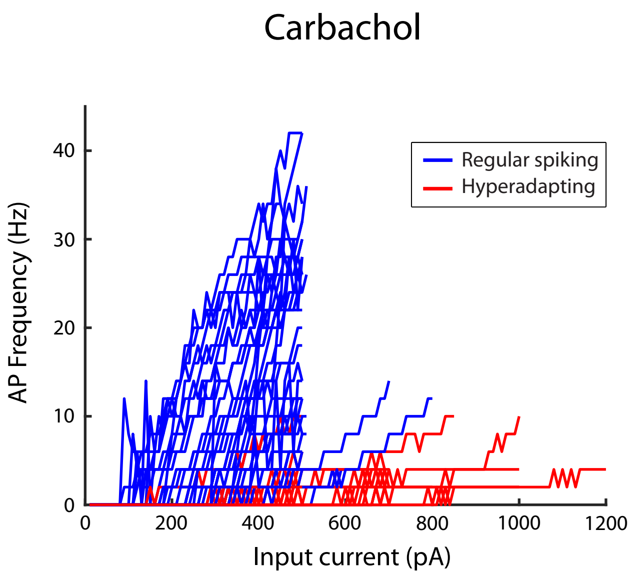
Carbachol
40
30
20
10
0
AP Frequency (Hz)
0
200
400
600
800
1000
1200
Input current (pA)
Regular spiking
Hyperadapting

**Figure S4**

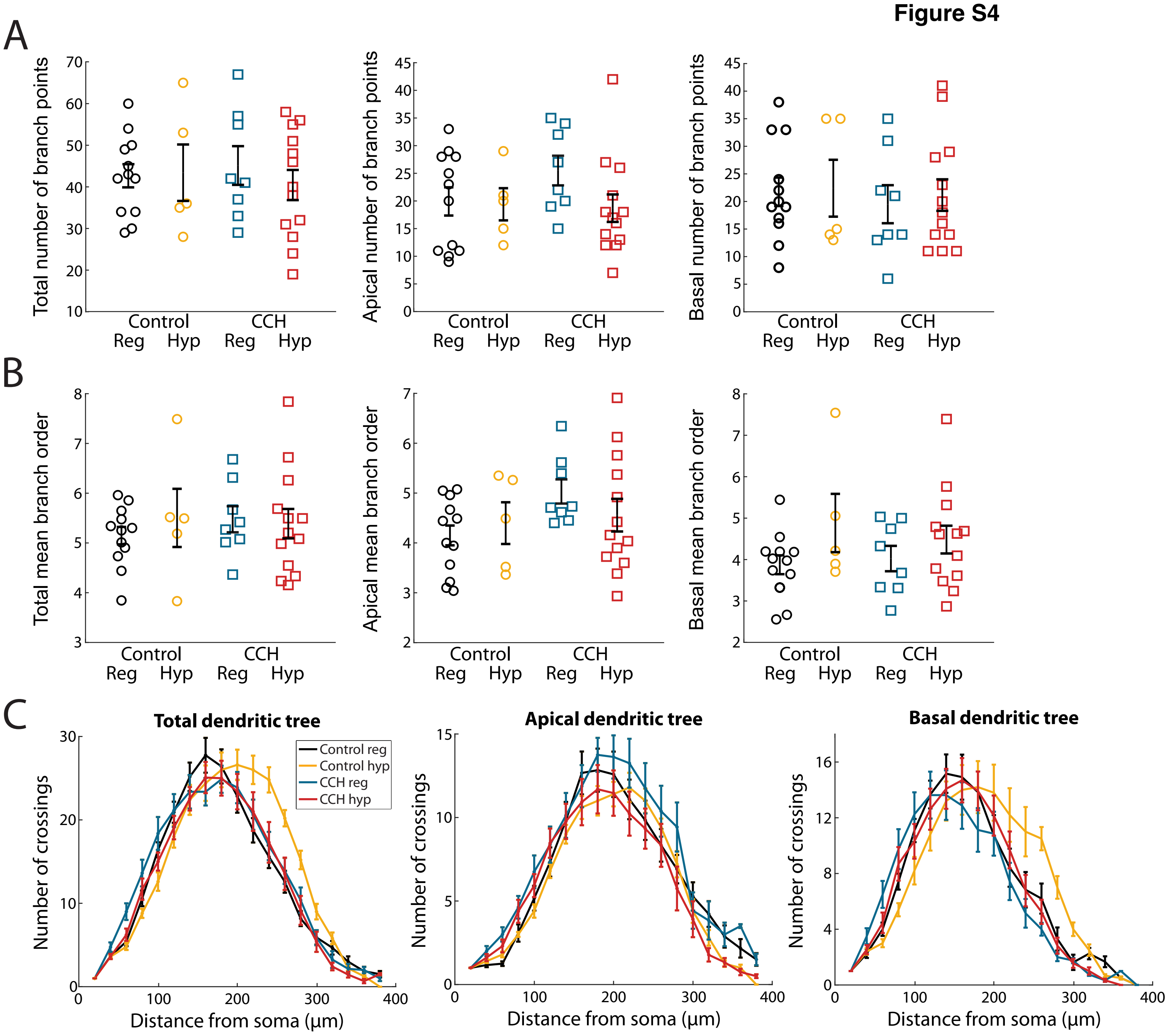
A
Total number of branch points
Apical number of branch points
Basal number of branch points
Control
CCH
Reg
Hyp
B
Total mean branch order
Apical mean branch order
Basal mean branch order
C
Total dendritic tree
Apical dendritic tree
Basal dendritic tree
Control reg
Control hyp
CCH reg
CCH hyp
Number of crossings
Distance from soma (µm)
